# Controlling lipid droplet dynamics via tether condensates

**DOI:** 10.1101/2024.05.15.594395

**Authors:** Chems Amari, Damien Simon, Theodore Bellon, Marie-Aude Plamont, Abdou Rachid Thiam, Zoher Gueroui

## Abstract

Lipid droplets (LDs) exhibit remarkable diversity and functionality within cells, depending on the metabolic needs of cells and the maintenance of lipid homeostasis. Such versatility is acquired through dynamic spatial and temporal positioning, enabling tight communication with other organelles. However, this complexity poses challenges in understanding LD biology. Controlled sequestration and release of LDs within their intracellular environment could offer a method to synchronize their behavior and better understand their function. Here, to advance in this direction, we developed ControLD (Controlled Trapping of Lipid Droplets), a novel approach designed to manipulate LDs and influence their dynamics and life cycle. By orchestrating the assembly/disassembly of engineered condensates, ControLD allows precise sequestration and release of LDs in cells. This technique effectively isolates LDs from the intracellular environment, drastically reducing interactions with other organelles. Notably, our experiments demonstrate that physically isolating LDs impairs their dynamics and remobilization during metabolic needs. ControLD represents a versatile tool for reversible LD trapping, with potential applications in controlling various cellular organelles.

## Introduction

Lipid droplets (LDs) are unique organelles at the nexus of cellular lipid metabolism. They consist of a neutral lipid core (mainly triacylglycerol, TG, and steryl ester) covered by a lipid monolayer embedded with proteins^1^. They play major roles in diverse biological functions, the primary of which is to balance cellular energy fluxes: storing excess energy and delivering lipids during scarcities, for making membranes or generating ATP^2,3^. In humans, dysregulation of LD formation and dynamics is related to several metabolic diseases linked to obesity, such as type II diabetes, liver or cardiac diseases, or lipodystrophies and neuronal disorders^2,4^. To perform their biological processes, LDs interact with nearly all other cell organelles, e.g. the endoplasmic reticulum (ER), mitochondria, autophagosomes, lysosomes, or peroxisomes, for anabolic or catabolic processes^5,6^. For instance, during metabolic shifts towards energy expenditure, clustered LDs dissociate, recruit at their surface enzymes such as lipases, e.g. ATGL and HSL, and use microtubules to associate with mitochondria to which they deliver fatty acids for ß-oxidation^7^. This exemplifies the relevance of LD spatial positioning relative to other organelles for its proper functioning^8,9^.

LD dynamics and functions are determined by proteins at the LD surface^10,11^. Most of these proteins come either from the cytosol, often associating with LDs by amphipathic helices, or from the ER membrane, associating with LDs with monotopic domains^10,11^. Several of the biophysical hallmarks of LD biogenesis and dynamics have emerged from *in vitro* reconstitution experiments^12,13^, or biochemically, following cell fractionation assays^14^, identifying key steps leading to LD nucleation, growth and stabilization^15^. Additionally, the development of techniques to characterize subcellular LD proteome and lipid composition reveals for example how changes in diet affect subcellular reorganization, with protein sequestration on LDs and contact remodeling with organelles^16–19^.

Despite these progresses, how LD biogenesis and functions are spatiotemporally controlled to coordinate response to metabolic need, stress, and cell fate, remains poorly understood^20^. The development of methodologies to probe and perturb LDs directly in cells could be instrumental in answering these questions^21^. Here, we opted for a strategy to develop intracellular compartments programmed to selectively confine LDs, by capitalizing on our ability to create biomolecular assemblies. Indeed, such an approach can be an efficient strategy to physically isolate and sequester selected structures from the rest of the cell, and eventually modify their biochemistry. Namely, our method, ControLD, i.e. Controlled Trapping of LDs, aimed at targeting and confining LDs in cells in a controllable and reversible manner. ControLD utilized engineered condensates as artificial compartments programmed to target specifically LDs to reorganize and sequester them in cells.

ControLD is based on the design of a chimeric protein scaffold, composed of a part that is prone to phase-separate in cells and a second one, composed of an LD targeting motif or protein, e.g., here, validated for Perilipin2 (Plin2). When produced in HeLa cells, our scaffolds eventually condensed at the surface of LDs to form micrometric condensates entrapping all LDs. These LD-containing condensates displayed the hallmarks of phase-separated condensates, including localized nucleation, growth, and coalescence of the condensed phases. Furthermore, we could trigger the release of LDs within a minute-time scale, by dissociating the condensates. LD-condensates robustly formed with 3 different LD-associated proteins, Plin1, Plin 2, Plin3, indicating that the LD-anchoring proteins could be used for a broad range of protein-LD affinities. LD contact sites with other organelles were likewise strongly reduced. Our study revealed that the condensates physically isolated LDs from the rest of the cytosol, thereby impacting LD dynamics such as impaired LD catabolism during metabolic needs.

## Results

### Bioengineered condensates programmed to interact with lipid droplets

To promote the interactions between LDs and engineered condensates, we developed a protein scaffold with two specific properties: the ability to undergo phase separation in the cytoplasm and the capability to interact with the surface of LDs (Fig. 1A). We selected 5Fm, known to undergo concentration-dependent and reversible phase separation^22,23^. As the LD protein targeting component, we opted for Perilipin2 (Plin2), a member of the perilipin family known to target LDs in most human cells^24,25^. As anticipated, the expression of 5Fm scaffolds lacking the LD surface protein (Plin2) for 24 hours in HeLa cells resulted in micrometric mCherry condensates scattered throughout the cytoplasm (Fig. 1B). These condensates did not interact with LDs labeled by emGFP-Plin2 (and stained with LipidTOX), indicating the absence of non-specific interactions between Plin2-labelled LDs and the 5Fm-based condensates (Fig. 1B). In contrast, when 5Fm-emGFP-Plin2 was co-expressed with mCherry-5Fm for 24 hours in cells, it led to the formation of large structures comprising LDs embedded within condensates (Fig. 1C-D). Typically, we observed approximately 2 to 6 large compartments per cell with a diameter of about 2-6 µm, encompassing tens of LDs (Fig. 1E). Importantly, all LDs present in HeLa cells were consistently trapped within the condensates (Fig. 1C-D). On closer examination, we observed that the 3D condensed phase typically separates individual LDs from each other (Fig. 2A). A second pattern of LD organization was observed when the stoichiometry of co-transfected emGFP-5Fm-Plin2/5Fm plasmids was increased from 1:20 to 1:4 (i.e., an increase in emGFP-5Fm-Plin2 quantity). Under these conditions, LDs were predominantly densely packed in a beads-on-a-string arrangement (Fig. 2B). While 5Fm-emGFP-Plin2 scaffolds were homogeneously localized around LDs, the mCherry-5Fm signal was enriched at the contacts between adjacent LDs, potentially forming optically unresolved condensates (Fig. S1).

**Figure 1.**
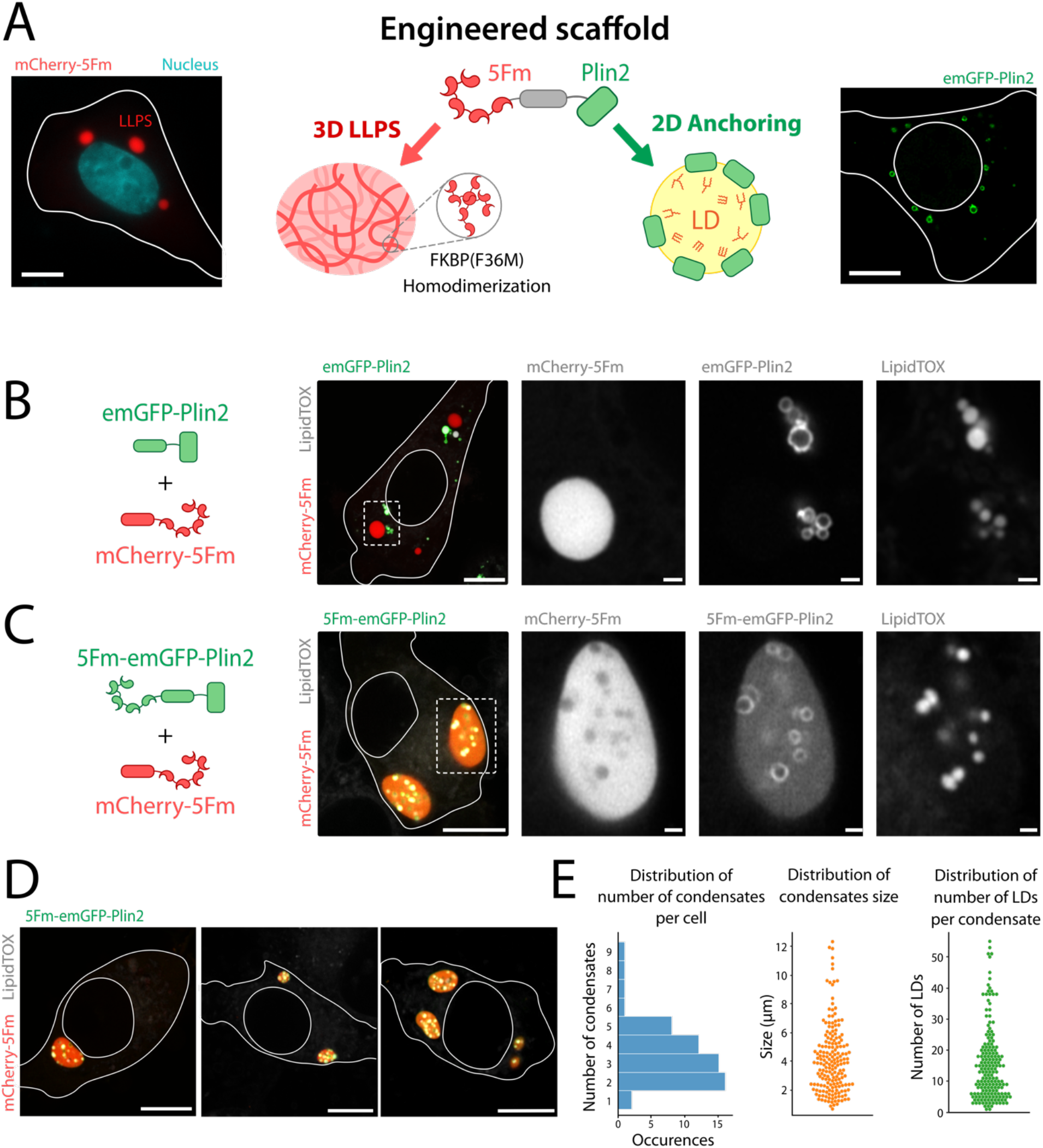
Controlled trapping of Lipid Droplets (ControLD) using bioengineered condensates programmed to interact with LDs. (A) Schematic of the LD-targeting LLPS protein scaffold. The scaffold is made of a repetition of homodimerization domains, and a LD-binding protein used for LD surface recognition (Fm = F36M-FKBP, Plin2 = perilipin 2). Left: HeLa cell expressing 5Fm condensates. Right: Hela cell expressing Plin2 localized at LD surface. Scale bar, 10 µm. (B) Airyscan confocal imaging of mCherry-5Fm condensates and Plin2-coated LDs in HeLa cells. Cells were fixed 24h after transfection with emGFP-Plin2, mCherry-5Fm, and 5Fm (plasmid ratio 1:5:14). LDs were stained with LipidTOX (white channel on merge). Grayscale images correspond to separate channels of the region of interest. Scale bar, 10 µm, 1 µm for zooms. (C) Airyscan confocal imaging of HeLa cells expressing Plin2-functionalized LLPS scaffolds. Cells were fixed 24h after transfection with 5Fm-emGFP-Plin2, mCherry-5Fm, and 5Fm (plasmid ratio 1:5:14). LDs were stained with LipidTOX (white channel on merge). Scale bar, 10 µm -, 1 µm for zooms. (D) Representative examples of cells expressing Plin2-functionalized LLPS scaffolds. Cells were fixed 24h after transfection with plasmids 5Fm-emGFP-Plin2, mCherry-5Fm, and 5Fm (plasmid ratio 1:5:14). Scale bar, 10 µm. (E) Left: distribution of the number of condensates per cell. Cells were fixed 24h after transfection with 5Fm-emGFP-Plin2, mCherry-5Fm, and 5Fm (plasmid ratio 1:5:14). Middle: size distribution of the condensates (Feret’s diameter). Right: distribution of the number of LDs per condensate (N = 52 cells from two independent experiments).

**Figure 2.**
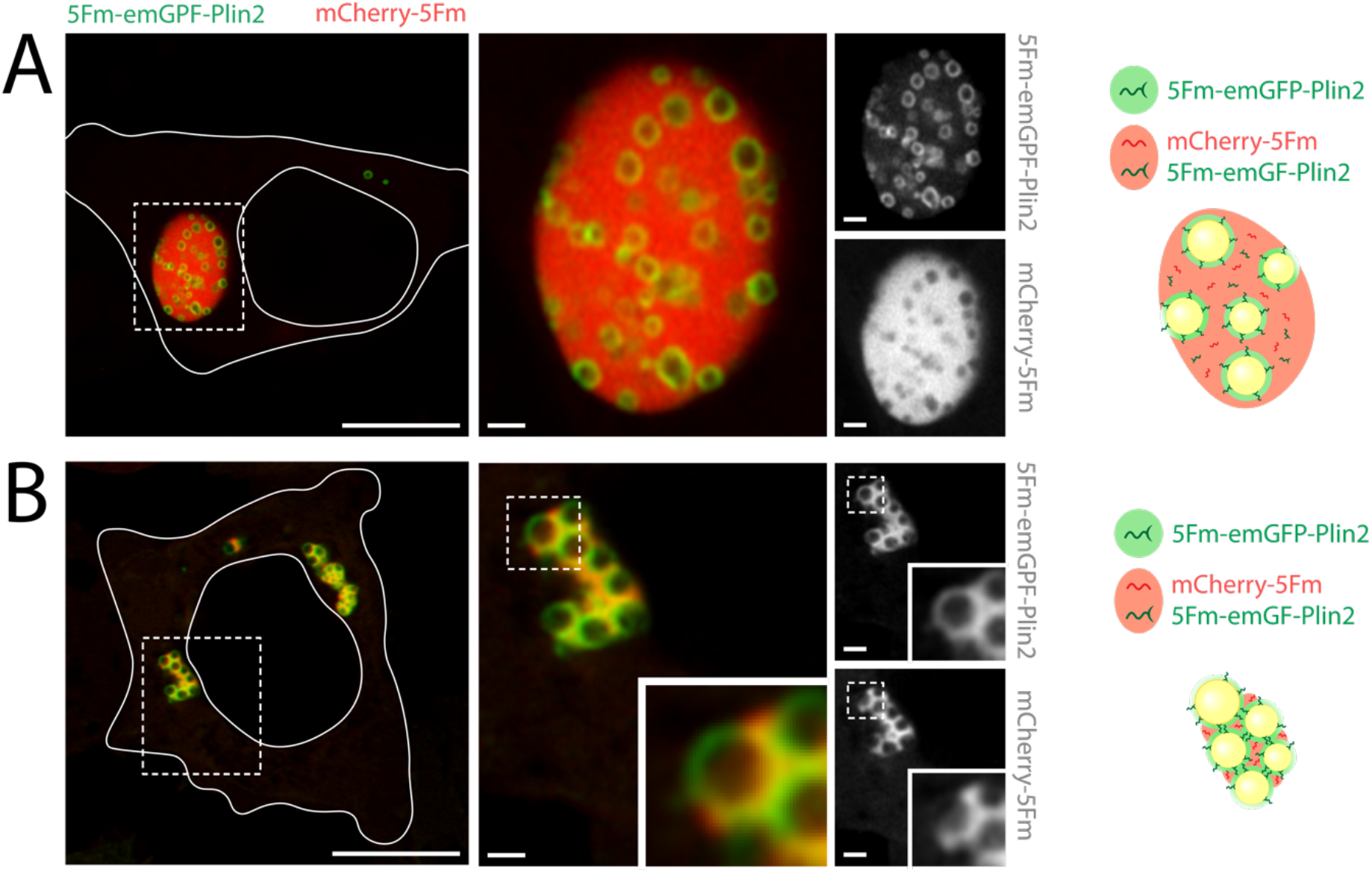
Morphology of LD-condensates. (A) Left: Airyscan confocal imaging of LD-condensates in HeLa cells observed at low Plin2 stoichiometry. Cells were fixed 24h after transfection with plasmids 5Fm-emGFP-Plin2, mCherry-5Fm, and 5Fm at low Plin2 stoichiometry (plasmid ratio 1:5:14). Scale bar, 10 µm, 1 µm for zooms. Right: schematic of the distribution of proteins composing LD-condensates. The protein corona depicted around LDs represents 5Fm-emGFP-Plin2, and the condensed phase symbolizes both protein scaffolds (5Fm-emGFP-Plin2 and mCherry-5Fm). (B) Left: Airyscan confocal imaging showing patterns of beads-on-a-string arrangements of LDs in HeLa cells at high Plin2 stoichiometry. Cells were fixed 24h after transfection with 5Fm-emGFP-Plin2, mCherry-5Fm, and 5Fm at high Plin2 stoichiometry (plasmid ratio 5:5:10). Scale bar, 10 µm, 1 µm for zooms. Right: schematic of the distribution of protein scaffolds.

Overall, our data demonstrate that bioengineered condensates programmed to target the LD surface can trap and confine LDs.

### LD surface triggers localized condensate formation

We next conducted live imaging to determine the timing and location of condensate formation within intracellular space. At early stages of protein expression, the 5Fm-emGFP-Plin2 signal localized at the surface of LDs, suggesting a 2D anchoring mechanism (Fig. 3A, S. Mov1). Subsequently, the mCherry-5Fm signal was recruited to the LD surface through Fm intermolecular interactions (Fig. 3A, S. Mov1). The condensates eventually nucleated on the LD surface for both 5Fm-emGFP-Plin2 and mCherry-5Fm scaffolds, indicating that LD surfaces played a role in seeding their growth. These LD-nucleated condensates tended to coalesce into larger structures, as commonly observed with classical liquid condensates (Fig. 3B). The typical time scale for the relaxation towards spherical bodies was approximately 30 minutes for a 5 µm-sized LD-condensate. Furthermore, we noted that LDs, initially in contact, moved apart during condensate growth (Fig. 3A, S. Mov1). Simultaneously with the initiation of LD-condensate formation, we observed that the cytosolic fluorescence intensity of emGFP-5Fm-Plin2 and mCherry-5Fm ceased increasing and reached a stable level whereas the total cytoplasmic fluorescence intensity (cytosolic + condensates) kept increasing over time (Fig. 3C, S2). This suggests that the nucleation of LD-condensates happens when a critical concentration of freely diffusing scaffolds is reached, as expected for phase separation driven by homotypic interactions. Overall, these results imply a sequential process in which the initial formation of a 2D scaffold layer takes place on the LD surface (as depicted in Fig. 3A), followed by the localized nucleation, growth, and coarsening of condensates, ultimately leading to the entrapment of LDs.

**Figure 3.**
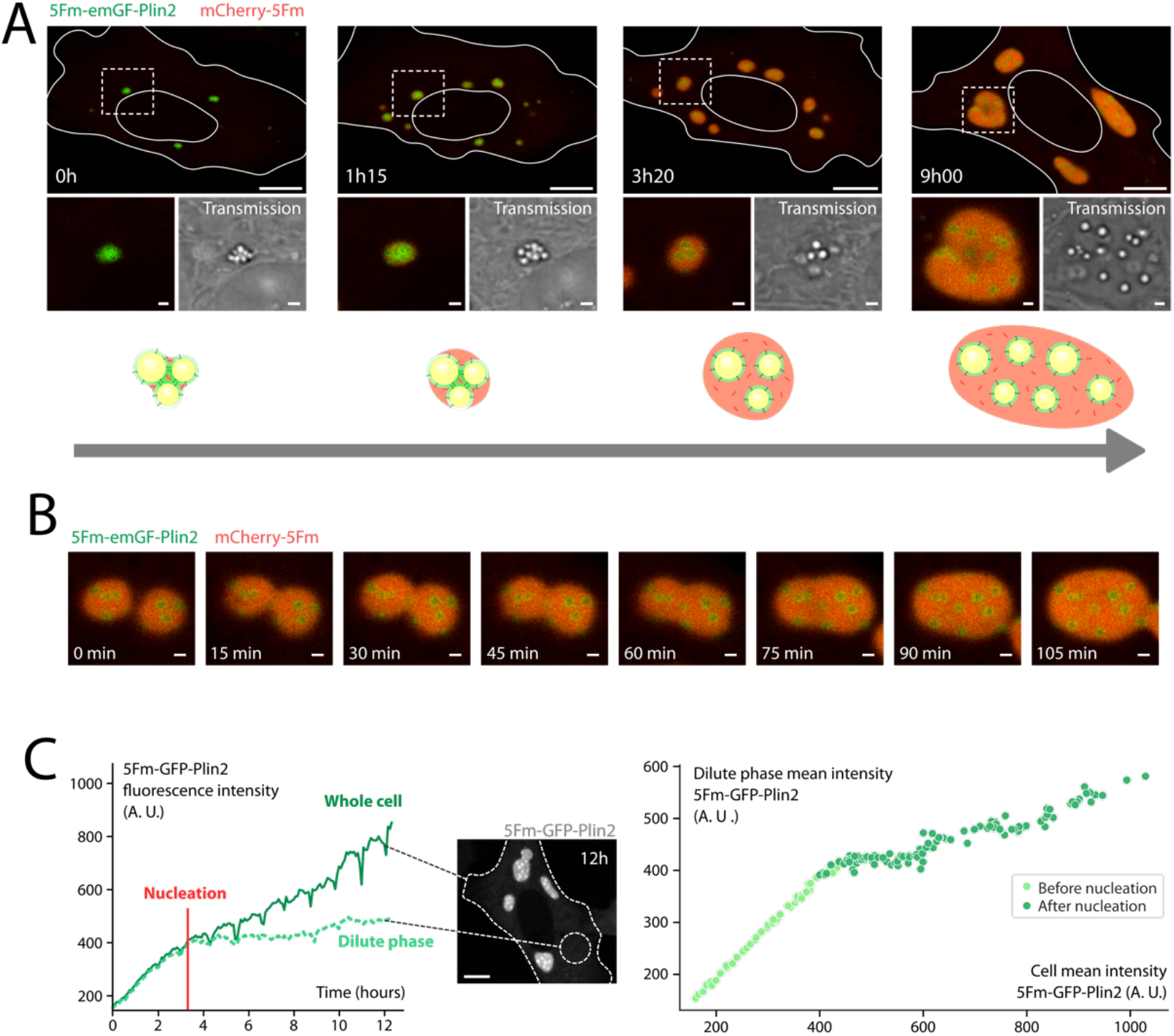
Dynamics of formation/disassembly of LD-condensates. (A) Time-lapse confocal imaging of the formation of LD-condensates in HeLa cells. Cells were transfected with 5Fm-emGFP-Plin2, mCherry-5Fm, and 5Fm (plasmid ratio 1:5:14), 6 hours before the start of the time-lapse. The region of interest shows the same condensates evolving during the time-lapse. The bottom panels display a zoom of the region of interest and a confocal transmission image of the same region. Scale bar, 10 µm, 1 µm for zooms. (B) Representative time-lapse confocal imaging showing the coalescence of two LD-condensates in HeLa cells, with the same transfection and observation conditions as for Fig. 3A. Scale bar, 1 µm. (C) Left: plot of the temporal evolution of the GFP mean fluorescence intensities during the formation of LD-condensates in a HeLa cell (transfection of 5Fm-emGFP-Plin2, mCherry-5Fm, and 5Fm; plasmid ratio 1:5:14). The overall intensity of the entire cell is associated with the mean intensity across the entire cell area, while the intensity in the dilute phase corresponds to the mean intensity in areas excluding LD-condensates. The “whole cell” intensity is associated with the mean intensity across the entire cell area, while the “dilute phase” intensity corresponds to the mean intensity in areas excluding LD-condensates. Right: evolution of solubilized 5Fm-emGFP-Plin2 fluorescence compared to total fluorescence, before and after condensates nucleation.

We next assessed whether the condensates could be disassembled by treating cells with FK506 (2.5 µM), a competitive binder that disrupts Fm-Fm interactions. Upon addition of FK506, we observed the rapid dissociation of the condensates, followed by the release of LDs, allowing them to move freely within the cytosol (Fig. 4, S.Mov. 2). This demonstrates the potential for targeted manipulation, entrapment, and release of LDs, thereby expanding the range of possibilities for druggable applications.

**Figure 4.**
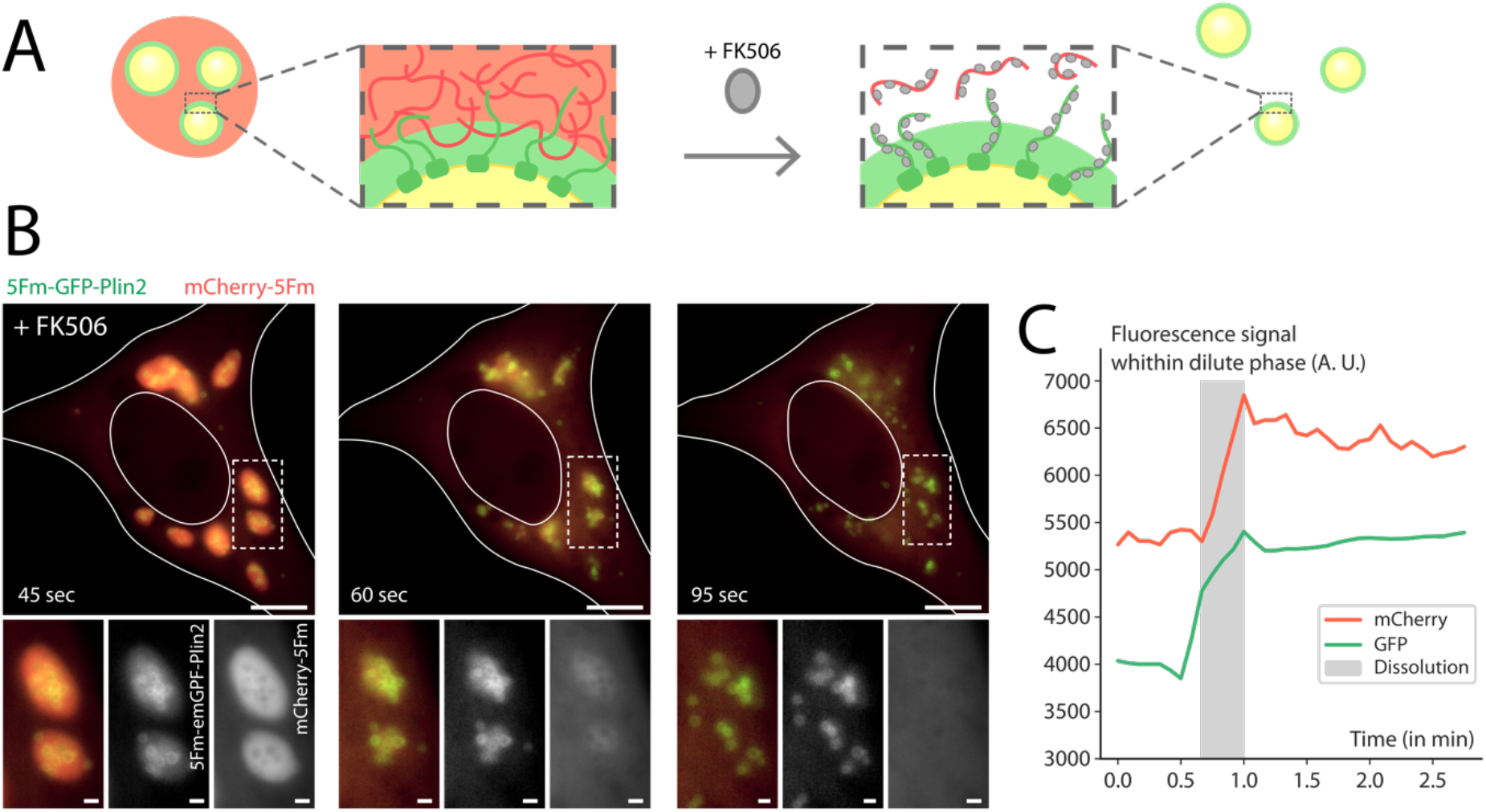
Drug-induced dissociation of LD-Condensates. (A) Schematic of FK506-induced dissolution of LD-Condensates. (B) Representative time-lapse epifluorescence imaging of the drug-induced dissolution of LD-trapping condensates in HeLa cells. Cells were transfected 24 hours before observation with 5Fm-emGFP-Plin2, mCherry-5Fm, and 5Fm (plasmid ratio 1:5:14). Dissolution is induced upon the addition of FK506 (2.5 µM) at the start of the time-lapse. Scale bar, 10 µm for full cell, 1 µm for zooms µm. (C) Evolution of fluorescence in the dilute phase during the condensate dissolution of the cell shown in B, for 5Fm-GFP-Plin2 and mCherry-5Fm. The grey area represents the visible dissolution of condensates.

### LD-condensates formation with Plin 1 and Plin 3 surface proteins

Plin2 belongs to the perilipin family of LD-associated proteins, known for their role in stabilizing LDs and regulating their surface accessibility^26^. To investigate the possibility of entrapping LDs using engineered condensates targeting alternative LD surface proteins, we focused on Plin 1 and Plin 3, which interact with stronger and weaker affinities, respectively, than Plin 2 to the LD surface (Fig. S3A-B)^27,28^. When co-expressing 5Fm-emGFP-Plin1 with mCherry-5Fm, we observed LD-condensates that exhibited a similar pattern to those observed for Plin2-LD-condensates (Fig. 5A). Specifically, LDs were embedded within condensates composed of 5Fm-emGFP-Plin1. Plin3-condensates were equally observed but only when we used a larger concentration of 5Fm-emGFP-Plin3 scaffolds (Fig. 5B). Together, these findings indicate the robust formation of LD-condensates with different LD-associated proteins characterized by distinct protein-LD affinities.

**Figure 5:**
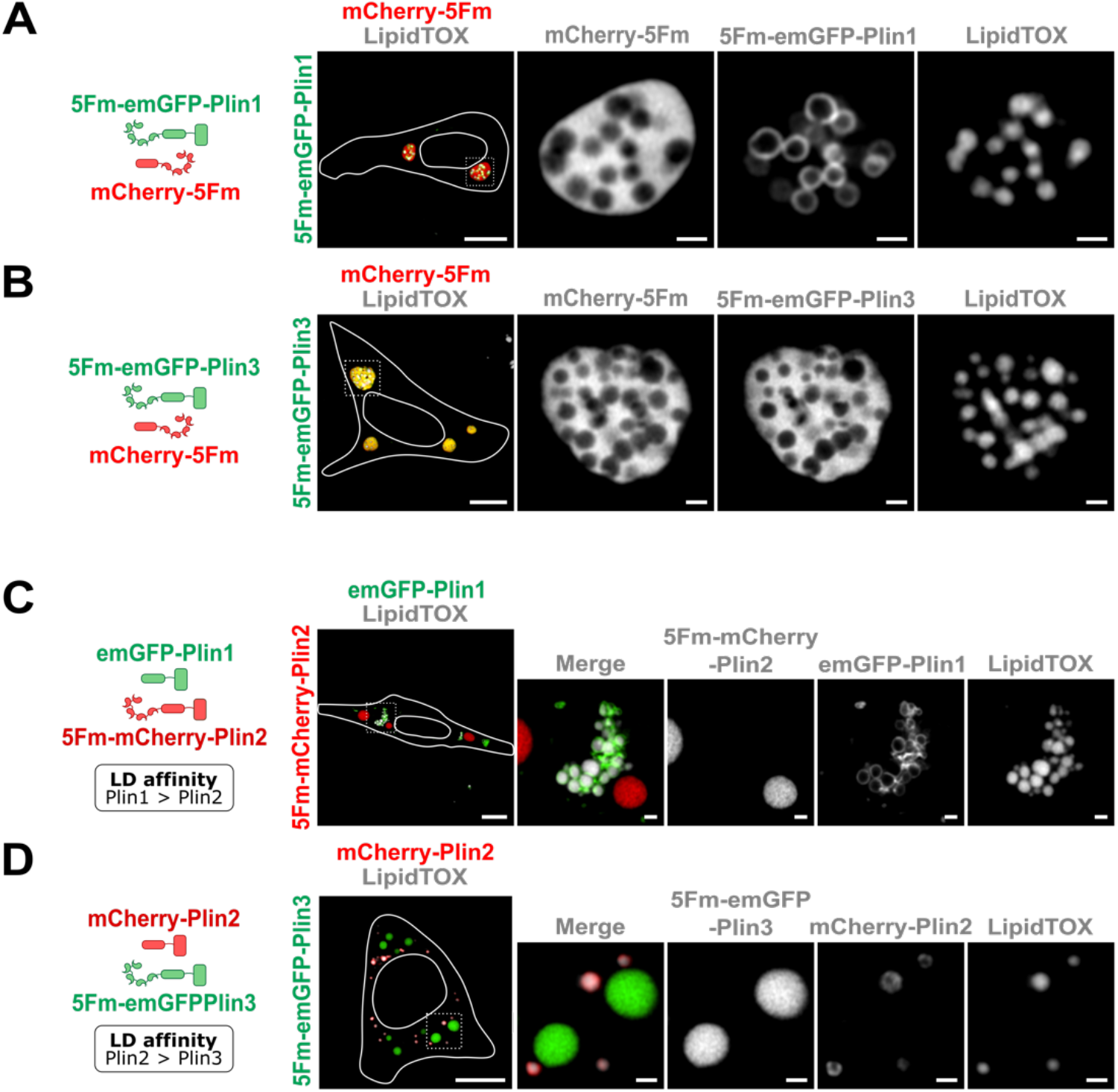
Formation of LD-condensates using Plin 1 and Plin 3 and protein binding competition. (A) Airyscan confocal imaging of LD-condensates in HeLa cells using Plin1 functionalized LLPS scaffolds. Cells were fixed 24 hours after transfection with plasmids 5Fm-emGFP-Plin1, mCherry-5Fm, and 5Fm (plasmid ratio 1:5:14). Scale bar, 10 µm, 1 µm for zooms. (B) Airyscan confocal imaging of LD-condensates in HeLa cells using Plin3 functionalized LLPS scaffolds. Cells were fixed 24 hours after transfection with plasmids 5Fm-emGFP-Plin3, mCherry-5Fm, and 5Fm (plasmid ratio 1:1:2). Scale bar, 10 µm, 1 µm for zooms. (C) Protein binding competition and condensate formation with cells co-transfected with emGFP-Plin1, 5Fm-mCherry-Plin2, 5Fm (plasmid ratio 1:1:19). (D) Protein binding competition and condensate formation with cells co-transfected with mCherry-Plin2, 5Fm-emGFP-Plin3, 5Fm (plasmid ratio 1:5:15). Scale bar, 10 µm, 1 µm for zooms. For all images, LDs were marked using LipidTOX™ HCS. Grayscale images correspond to separate channels of the region of interest.

### Protein binding competition at the LD surface impacts condensate formation

Since LD proteome is regulated by protein binding competitions occurring at the LD monolayer surface^29^, we probed the impact of protein competition on condensate formation. Knowing that Plin1 exhibits a stronger affinity with LDs than Plin2^27,28^, we assessed the effect of such competition on LD-condensate formation by co-transfecting Plin2 condensate scaffolds (5Fm-emGFP-Plin2 and 5Fm) with emGFP-Plin1. Whereas emGFP-Plin1 strongly labeled LD surface as expected, Plin2-condensates were devoid of LDs (Fig. 5C). This behavior was also observed when co-transfecting Plin3 condensate scaffolds with mCherry-Plin2, for which we found that mCherry-Plin2 were labeling all LDs and Plin3-condensates formed without trapping any LDs (Fig. 5D). The saturation of the LD surface with a higher-affinity protein could prevent the targeting of LD-functionalized LLPS scaffolds and consequently the entrapping of LDs. Therefore, the over-expression of Plin of stronger affinity, which binds preferentially to the LD surface, resulted in the formation of condensates of Plin of weaker affinities that are devoid of LDs.

### LD confinement restricts inter-organelle interactions

LD-organelle contacts are central to many processes involving material transfer, signaling, and lipid metabolism^30^. The observed perturbation of LD dynamics using ControLD supports the hypothesis that LD confinement could disrupt LD-organelle interactions and contact sites. We investigated the spatial organization of confined LDs with ER and mitochondria to verify this hypothesis. We first examined Plin1 which relocates from the ER to LDs^24,27,31^, and found that LDs labeled by emGFP-Plin1 were juxtaposed to ER structures labeled with mCherry-Sec61 (Fig. 6A). Interestingly, when expressing 5Fm-emGFP-Plin1, we found that ER structures were excluded from Plin1-condensates, indicating in this case a strong restriction of LD-ER inter-organelle interactions. This absence of LD-ER contacts was also observed using Plin2- and Plin3-condensates (Fig. 6B, S4A). Next, we visualized mitochondria and LDs in cells expressing Plin-condensates and stained with MitoTracker DeepRed (Fig. 6C, S4B). Likewise, we found that mitochondria structures were generally excluded from Plin-condensates for all three Plin-type condensates, indicating an absence of contacts with confined LDs. These findings suggest that the tether condensates restrict inter-organelle interactions.

**Figure 6:**
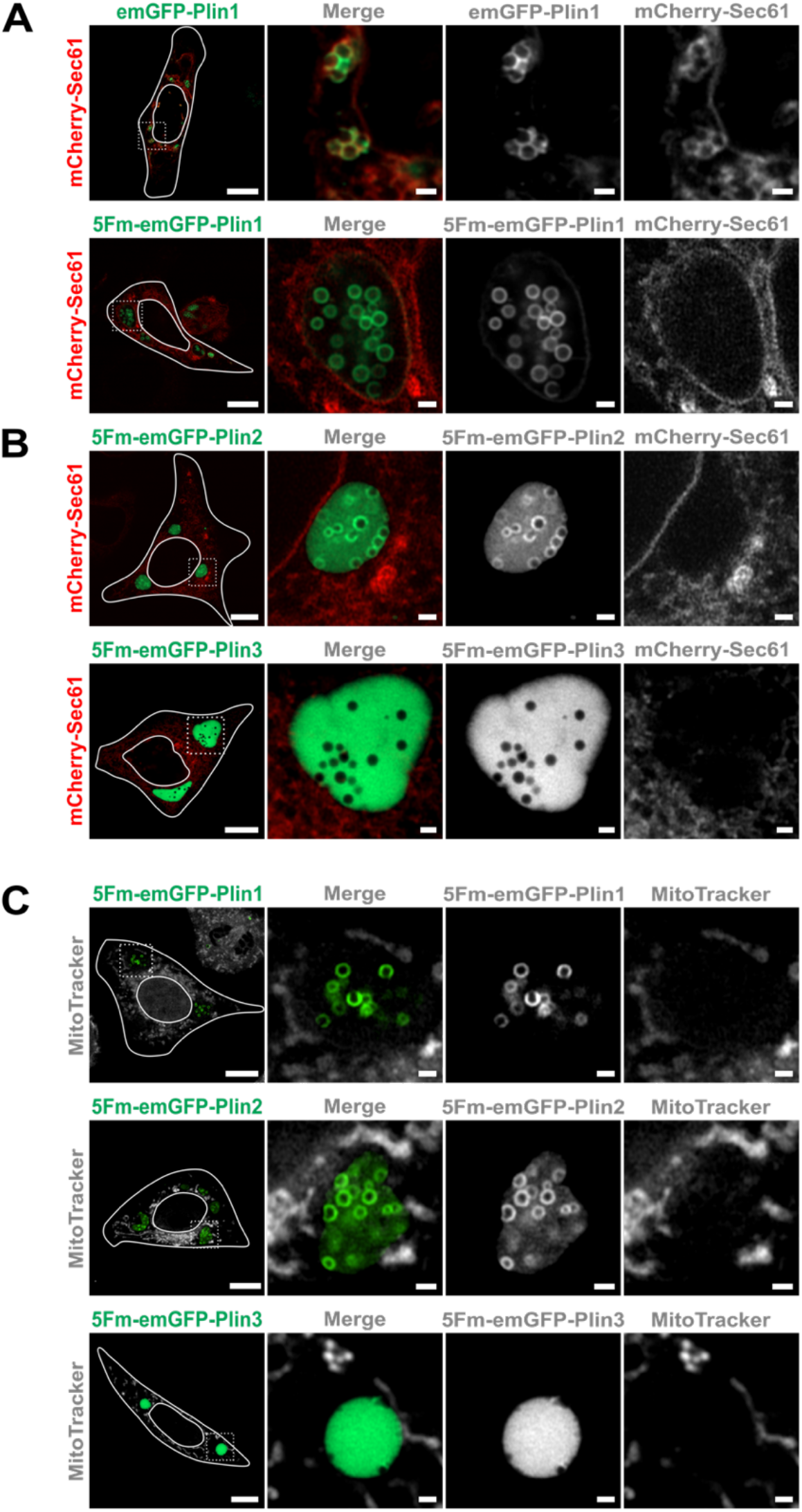
LD-condensates interactions with ER and mitochondria. (A) Representative confocal imaging of LD-condensates in HeLa expressing for 24 hours emGFP-Plin1 with mCherry-Sec61 (top panel) or 5Fm-emGFP-Plin1 with mCherry-Sec61 (lower panel). Scale bar, 10 µm and 1 µm for zooms. (B) Confocal imaging of LD-condensates in HeLa expressing for 24 hours 5Fm-emGFP-mPlin2 with mCherry-Sec61 (top panel) or 5Fm-emGFP-mPlin3 with mCherry-Sec61 (lower panel). Scale bar, 10 µm and 1 µm for zooms. (C) Confocal imaging of LD-condensates in HeLa expressing for 24 hours 5Fm-emGFP-Plin1 or 5Fm-emGFP-Plin2 or 5Fm-emGFP-Plin3, in the presence of a mitochondria marker. Mitochondria were revealed using MitoTracker™ Deep Red FM. Scale bar, 10 µm and 1 µm for zooms.

### Effect of LD confinement on LD dynamics

The confinement of LDs mediated by condensates may act as a shield limiting LD surface accessibility, which in turn could have implications for their life cycle and dynamics. This confinement could jeopardize LD function as storing energy sources when cells encounter fluctuations in nutrients. To investigate this possibility, we studied the consumption of LDs, confined or not, during cell starvation. The induction of starvation was realized by changing the cell culture media to a nutrient-low salt solution (EBSS). As expected, we observed that unconfined LDs were entirely consumed when cells were placed in the nutrient-deprived media for 24 hours, in contrast to cells placed in complete media as shown by Plin2-coated signals (Fig. 7A). Most interestingly, we observed a large number of LDs within condensates in the starved cells, indicating that they were protected from consumption by the confinement (Fig. 7B). To further analyze the effect, we quantified the total number of LDs in cells in an unconfined or confined situation, i.e, either expressing 5Fm condensates or LD-condensates, both with and without the induction of starvation (Fig. 7). We found a 7-fold decrease of the mean of LD numbers in cells by inducing starvation for the cells displaying unconfined LDs (49 LDs/cell to 7 LDs per cell, mean numbers). In comparison, this decrease was only about 1.2 folds for the cells containing LD-condensates (35 LDs/cell to 27 LDs/cells, mean numbers). Altogether, this reveals that condensates are impeding the metabolic consumption of contained LDs during cell starvation, most probably due to a restriction of LD surface accessibility.

**Figure 7.**
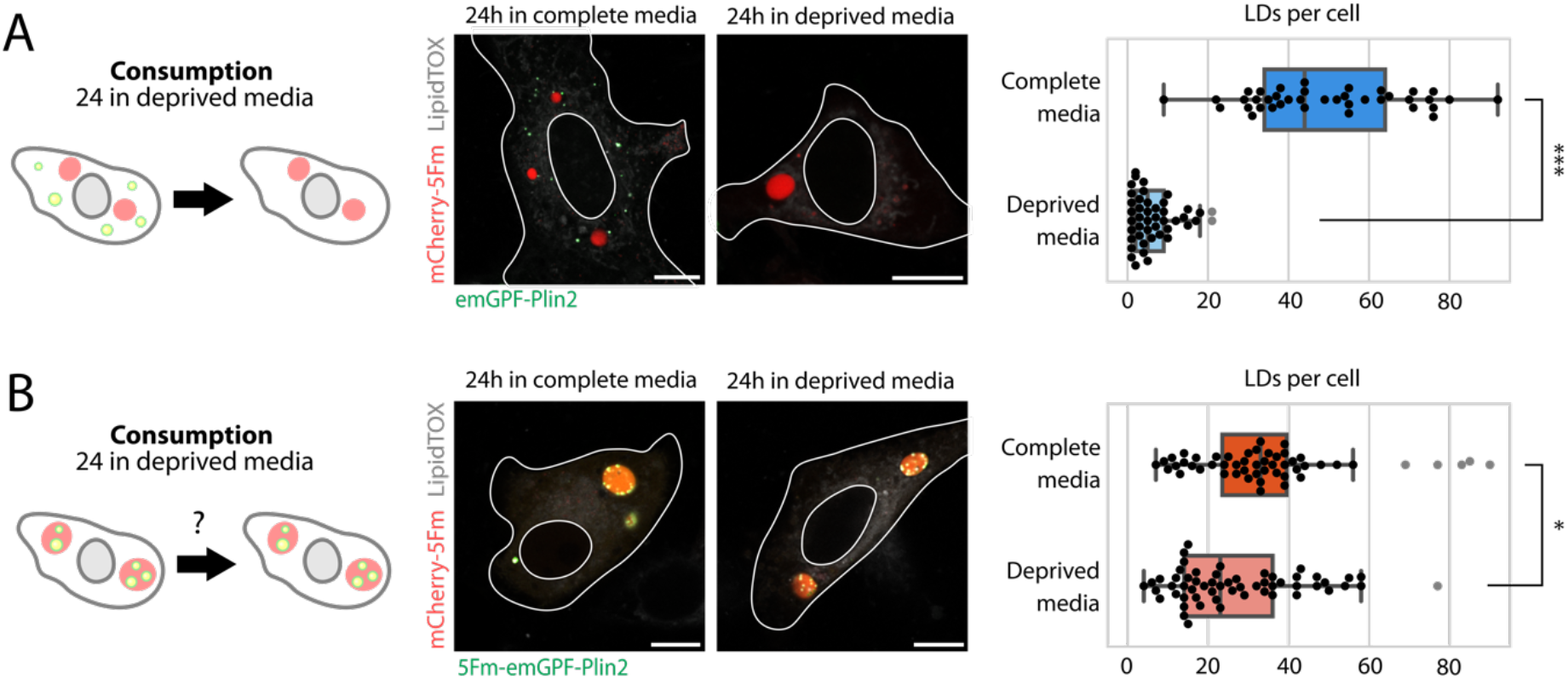
LD-trapping condensates protect LDs from consumption during cell starvation. (A) Effect of starvation on unconfined LDs. Center: epifluorescence imaging of Plin2-coated LDs and 5Fm-mCherry condensates in HeLa cells, 48 hours after transfection with plasmids 5Fm-emGFP-Plin2, mCherry-5Fm, and 5Fm (plasmid ratio 1:5:14). Culture media was changed 24 hours before fixation, either by renewing the media (complete media), or by switching to a deprived media (EBSS + 1% BSA). Scale bar, 10 µm. Right: Mean number of LDs per cell, for cells expressing at least one 5Fm condensate, with and without starvation. (p-value: 8.10^-15^). (B) Effect of starvation on confined LDs. Right: epifluorescence imaging of LDs confined in condensates in HeLa cells, 48 hours after transfection with plasmids 5Fm-emGFP-Plin2, mCherry-5Fm, and 5Fm (plasmid ratio 1:5:14). Culture media was changed 24 hours before fixation, either by renewing the media (complete media), or by switching to a deprived media (EBSS + 1% BSA). Scale bar, 10 µm. Right: Mean number of LDs per cell, for cells expressing at least one LD-Condensate, with and without starvation. (p-value: 4.3·10^-3^).

## Discussion

We developed ControLD, a versatile method to trap and confine LDs within engineered condensates. To demonstrate its effectiveness, ControLD was tested using Perilipin-functionalized phase-separating scaffolds. These scaffolds specifically target the surface of LDs and form a condensed protein meshwork, effectively isolating all LDs from the surrounding intracellular environment. The LD trapping is reversible since LDs can be released by triggering the disassembly of the condensates within a minute timescale. Using ControLD, we demonstrated that the physical isolation of LDs by the condensates drastically reduces their interactions with mitochondria and the endoplasmic reticulum, and most likely with other organelles. Importantly, we showed that by physically isolating LDs from the rest of the cytosol, LD dynamics and remobilization during metabolic needs were impaired.

The life cycle and dynamics of LDs are central to the metabolic roles they play, and such processes are tightly linked to specific biochemical mechanisms occurring at LDs’ surface and LD-organelle contact sites. How does entrapping LDs within condensates result in arresting such LD dynamics? We could envision distinct but not exclusive mechanisms. Firstly, condensates may act as a diffusion barrier, preventing enzymes such as lipases from targeting the LD surface. Secondly, the condensate physical barrier could limit the initiation of the degradation machinery of LDs, e.g. mediated by lipophagy or lysosomes^32^. Thirdly, contact sites between LDs and other organelles could be strongly perturbed, leading to altered communication between organelles. All these mechanisms could play collectively, enabling ControLD to manipulate LD metabolism and interactions with organelles reversibly.

Beyond its technical characteristics, ControLD opens up avenues for exploring the significance of condensates in regulating LD biology. While mounting evidence shows an interplay between membrane-bound organelles, e.g., ER, Golgi, plasma membrane, mitochondria, and condensates, there is currently no conclusive evidence that LDs might interact with condensates. Yet, a description of the interplay between LD dynamics and protein phase separation could provide a fresh perspective on regulatory mechanisms to tune the LD life cycle. For instance, it has been proposed that the condensation of Cidec proteins at LD-LD contact sites could facilitate LD fusions^33^. Interestingly, our condensates mimic phase-separated systems in cells and could provide a framework for a better understanding of how LDs and phase-separated condensates may interact.

In our study, we found that the nucleation of condensates occurred on LDs by a step-wise formation process with the recruitment of LLPS scaffolds at the LD surface, followed by their localized condensation (Fig. 3). LD surface provides numerous anchoring sites for disordered motifs such as amphipathic helices^34^. Therefore, LDs may favorably recruit LLPS scaffolds and restrict their mobility on 2D in comparison to free diffusing in the cytosol. This restriction in scaffold diffusion could reduce the concentration threshold for condensation, as proposed for the regulation of phase-separated condensates by membranes^35^. Due to its higher phospholipid packing voids than bilayer membranes^36^, it is perfectly plausible that the LD surface facilitates more LLPS nucleation events.

Our observations of condensate nucleation triggered at the LD surface are also consistent with *in vitro* experiments using the histidine-modified FUS low complexity domain interacting with NTA(Ni)-phospholipids^37^. In our system, the mechanism of recruitment of LLPS scaffolds on the LD surface is mediated by Plin proteins, which are LD-specific proteins. Other biomolecules than LD surface proteins are likely to bind to their surface, including RNA or RNA Binding Proteins ^38^, which could act as putative linkers with endogenous condensates. Our data suggest that the surface of LD could drive the nucleation of condensates, which implies a general mechanism of regulation of phase separation that is based both on the biophysical and biochemical properties of lipid droplet/condensate interactions.

Physical considerations may also explain how LD organizations targeted with Plin-functionalized scaffolds could be remodeled through two main different patterns, either entrapped within large condensates or arranged into a string-like pattern (Fig. 2). Interestingly, the two patterns were found to depend on Plin2-5Fm expressed in cells, with a lower quantity of Plin2 favoring large 3D condensates. Our time-lapse imaging suggests that LDs-on-a-string arrangements formed through the irreversible adhesion between close LDs (Fig. 2B). In these conditions, large condensates on LD surface failed to grow. These patterns were mainly observed when a larger amount of Plin2-5Fm was transfected in comparison to the quantity used for forming LD-condensates (five-fold increase). This indicates that competition between adhesion strength, dependent on the Plin2-5Fm surface density, and 3D phase separation dictates the balance between LDs-on-a-string arrangements and LD-condensates. This mechanism could potentially elucidate why, under physiological conditions, LDs often cluster resembling a grape-like structure, potentially facilitated by proteins prone to forming condensates. When metabolic demands arise, these adhesion interactions may be disrupted through the removal of such proteins, allowing for LDs to disperse and fulfill their metabolic functions. Lastly, we anticipate that the concept of ControLD, which allows for the temporal control of trapping/releasing of LDs, could be applied to diverse applications, including the study of LD biology to its extension for controlling other cellular organelles.

## METHODS

### Experimental model

Human epithelioid carcinoma HeLa cells (ATCC, ccl‐2) were kept in Dulbecco’s modified Eagle’s medium (with 4.5 g/l D‐glucose, HyClone) with 10% fetal bovine serum (Gibco) and 1% penicillin–streptomycin (Sigma, P4333), at 37°C in a 5% CO2‐humidified atmosphere. Tests for mycoplasma contamination were routinely carried out.

### Plasmids

The plasmids used during this study are listed in the table S1. The pcDNA3.1-5Fm and pcDNa3.1-mCherry-5Fm plasmids were obtained as described in^23^. The pcDNA3.1-emGFP-mPlin2 plasmid was created by replacing the 5Fm coding sequence from the pcDNA3.1-emGFP-5Fm (between the Eco47III and XbaI restriction sites) with the Plin2 coding sequence. The pcDNA3.1-5Fm-emGFP-Plin2 plasmid was created by inserting the mPlin2 coding sequence between the XbaI and AgeI restriction sites from the pcDNA3.1-5Fm-emGFP. The pcDNA3.1-5Fm-emGFP-Plin1 plasmid was created by replacing the Plin2 coding sequence from the pcDNA3.1-5Fm-emGFP-mPlin2 (between the XbaI and AgeI restriction sites) with the Plin1 coding sequence. The pcDNA3.1-5Fm-emGFP-Plin3 plasmid was created by replacing the mPlin2 coding sequence from the pcDNA3.1-5Fm-emGFP-Plin2 (between the XbaI and AgeI restriction sites) with the mPlin3 coding sequence. The pcDNA3.1-emGFP-Plin1 plasmid was created by replacing the mPlin2 coding sequence from the pcDNA3.1-emGFP-Plin2 (between the AfeI and XbaI restriction sites) with the mPlin1 coding sequence. The pcDNA3.1-5Fm-mCherry-Plin2 plasmid was created by replacing the emGFP coding sequence from the pcDNA3.1-5Fm-emGFP-Plin2 (between the SacII and XbaI restriction sites) with the mCherry coding sequence. The pcDNA3.1-mCherry-Plin2 plasmid was created by replacing the 5Fm coding sequence from the pcDNA3.1-mCherry-5Fm (between the Eco47III and XbaI restriction sites) with the Plin2 coding sequence. The pcDNA3.1-emGFP-Plin3 plasmid was created by replacing the 5Fm coding sequence from the pcDNA3.1-emGFP-5Fm (between the Eco47III and XbaI restriction sites) with the mPlin3 coding sequence.

### Transfection

For imaging after cell transfection and fixation, HeLa cells were cultured on 22×22 mm glass coverslips (VWR) in 6‐well plates (Falcon, 3.5 × 10^5^ cells per well). For live imaging, Hela cells were seeded on 35‐mm dishes with a polymer coverslip bottom (Ibidi, 1.5 × 10^5^ cells per dish). In both cases, adhesion surfaces were treated with a solution of 0.002% poly-L-lysine (Sigma) for 30 min. Cells were transfected with Lipofectamine 2000 (Invitrogen), using 2 µg of DNA, 24 hours after seeding. For fixed cell imaging, cells were transfected with plasmids and ratios as indicated in the legends. In every case, cells were incubated at 37°C under a 5% CO_2_ atmosphere.

To prepare fixed samples, cells were washed two times with PBS, and then incubated 20 min in a solution of 4% PFA (polyformaldehyde, Fisher Scientific) in PBS. Cells were washed again two times with PBS, and mounted on slides (VWR) using Vectashield mounting medium (Vector Laboratories, H‐1200).

### Live dissolution assay

To induce the live dissolution of condensates (Fig. 5, Supplementary Movie 2), a drop of FK506 (Sigma, F4679) at 2.5 mM in DMSO was added in the medium at the edge of the culture plate, after the start of live acquisition. No mixing was performed to allow the slow diffusion of FK506 in the medium, to reach a final concentration of 2.5 µM.

### Delipidation assays

To induce the delipidation of cells, HeLa cells were first seeded and transfected 24h later as previously described, on a 22×22 mm coverslip in a 6-well plate. After another 24h, cells were washed 2 times with pre-heated DPBS (Dulbecco Phosphate-Buffered Saline, Sigma) and placed in EBSS (Earle’s Balanced Saline Solution, Fisher Scientific) containing 1% of BSA (Bovine Serum Albumine, Sigma) during 24h. Control cells were instead placed again in the same culture medium as the first steps during 24h. Cells were then fixed and observed as described above.

### Imaging

For overnight live imaging (Fig. 3, Supplementary Movie 1), cells were imaged on a Zeiss LSM 710 confocal microscope using an x63 oil-immersion objective (PlanApochromatic, numerical aperture (NA) 1.4), at 37°C under a 5% CO_2_ atmosphere. The acquisition was started 6h hours after transfection for the formation of condensates. Microscope hardware and image acquisition were controlled with LSM Software Zen 2012. For the dissolution assay, we used (Fig. 5, Supplementary Movie 2), cells were imaged on an Olympus IX81 microscope and 63× oil immersion objective (PlanApo, NA 1.42), equipped with a CMOS camera, Orca‐Fusion (Hamamatsu, Corporation), and a LED system of illumination (Spectra X, Lumencor) at 37°C. The acquisition was started 24h hours after cell transfection. Microscope hardware and image acquisition were controlled using Micro-Manager. For Airyscan images, cells were imaged with Zeiss LSM 880 confocal microscope equipped with Airyscan detector using a 60× oil immersion objective (PlanApo, NA 1.42). Images were Airyscan-processed automatically with Zeiss Zen software (v. 2.3).

### Data analysis

To measure mean fluorescence signals in cells (Fig. 3C, Fig. S2), cells were systematically segmented using Cellpose^39,40^ using a pre-trained model which was fine-tuned on our data, across an overnight time-lapse movie. Fluorescence mean values were then computed using Python. To obtain the signal in the cytosol only, condensates were systematically thresholded using Otsu threshold^41^ provided by the scikit-image Python package^42^, and fluorescence mean values were computed on the remaining areas of the cell as before.

Plotted fluorescence profiles (Fig. S1) were generated using Fiji^43^. All plots were generated using the seaborn Python package^44^.

### Statistical analysis

To measure the distribution of condensate features (Fig. 1E), 52 cells expressing condensates from two independent experiments were used. Counting was done manually, and size measurement was performed using Fiji.

Counting of LDs in cells during the delipidation experiments (Fig. 7C) was performed by blind 3D acquisitions of at least 40 cells expressing condensates for each condition, from two independent experiments, followed by manual counting of all LDs in the 3D stack. P-values were calculated using a two-sided Wilcoxon-Mann-Whitney provided by the Scipy Python package^45^.

All statistical analyses were performed using Python and plotted using Seaborn. Regarding the plots of Fig. 1, 52 cells from two independent experiments were observed through blind acquisition and analyzed. Regarding the boxplots of Fig. 7, the middle line of the boxplots displays the median value, and its width symbolizes the inner quartiles. The number of cells for these plots were respectively 39 (unconfined LDs and complete medium), 45 (unconfined LDs and starving medium), 52 (confined LDs and complete medium), and 57 (confined LDs and starving medium), and cells from two independent experiments were observed though blind acquisition, by selecting every cell expressing a condensate.

## Acknowledgments and Funding

We thank Mohyeddine Omrane for the help along the project. CA was supported by a PhD fellowship from ENS. D.S was supported by the Labex IPGG (ANR-10-IDEX-0001-02). This work was supported by the centre national de la recherche scientifique, Ecole Normale Superieure, and ANR (ANR-22-CE44-0036) to A.R.T and Z.G, and foundation ARC(ARCPJA2023080007011) to Z.G.

## Author Contributions

Chems Amari: Conceptualization; Investigation; data curation; methodology; writing – original draft; writing – review and editing.

Damien Simon: Investigation; data curation; methodology; writing – original draft. Théodore Bellon: Investigation; methodology.

Marie Aude Plamont: Investigation; methodology.

Abdu Rachid Thiam: Conceptualization; methodology; resources; funding acquisition; writing – review and editing.

Zoher Gueroui: Conceptualization; methodology; resources; supervision; data curation; funding acquisition; writing – original draft; writing – review and editing.

## List of Supplementary Materials

4 Supplementary Figures, 1 Supplementary Table, 2 Supplementary Movies.

## Data and materials availability

All data needed to evaluate the conclusions in the paper are present in the paper and/or the Supplementary Materials.

