## Supplementary Files for "Controlling lipid droplet dynamics via tether condensates"

**Supplementary Table 1**
**4 Supplementary Figures**
**Legend for Supplementary Movies**

### Supplementary Table 1

| Plasmid | Source | Fwd Primer Sequence(s) | Rev Primer Sequence(s) |
|---|---|---|---|
| pcDNA3.1-mCherry-5Fm | Cochard *et. al.* 2023 | n/a | n/a |
| pcDNA3.1-emGFP-mPlin2 | This article | GTCAGTAGCGCTGCAGCAGCAGTAGTGGATC | ACTGACTCTAGACTGAGCTTTGACCTCAGAC |
| pcDNA3.1-5Fm-emGFP-mPlin2 | This article | CTCGAGTCTAGAGCATCCGTTGCAGTT | CTCGATACCGGTCAGATCCTCTTCTGAGATGAGTTTTTGTTCGAATTCATGAGTTTTA |
| pcDNA3.1-5Fm-emGFP-mPlin1 | This article | gctagctctagaATGTCAATGAACAAGGGCCCAACC | gctagcaccggtcagatcctcttctgagatgagtttttgttcgaattcGCTCTTCTTGCGCAGCTGG |
| pcDNA3.1-5Fm-emGFP-mPlin3 | This article | gtcagttctagaATGTCTAGCAATGGTACAGATGCG | gtcagtaccggtcagatcctcttctgagatgagtttttgttcgaattcCTTCCCTTCAGGGGTTTTCTC |
| pcDNA3.1-emGFP-mPlin1 | This article | gctagcAGCGCTATGTCAATGAACAAGGGCCCAACC | gctagcTCTAGAGCTCTTCTTGCGCAGCTGG |
| pcDNA3.1-5Fm-mCherry-mPlin2 | This article | restriction enzyme used (SacII) with pcDNA3.1-5Fm-emGFP-mPlin2 | restriction enzyme used (SacII) with pcDNA3.1-5Fm-emGFP-mPlin2 |
| pcDNA3.1-mCherry-mPlin2 | This article | restriction enzyme used (SacII) with pcDNA3.1-mCherry-5Fm | restriction enzyme used (SacII) with pcDNA3.1-mCherry-5Fm |
| pcDNA3.1-emGFP-mPlin3 | This article | gtcagtagcgctATGTCTAGCAATGGTACAGATGCG | actgactctagaCTTCCCTTCAGGGGTTTTCTC |

**Supplementary Figures**

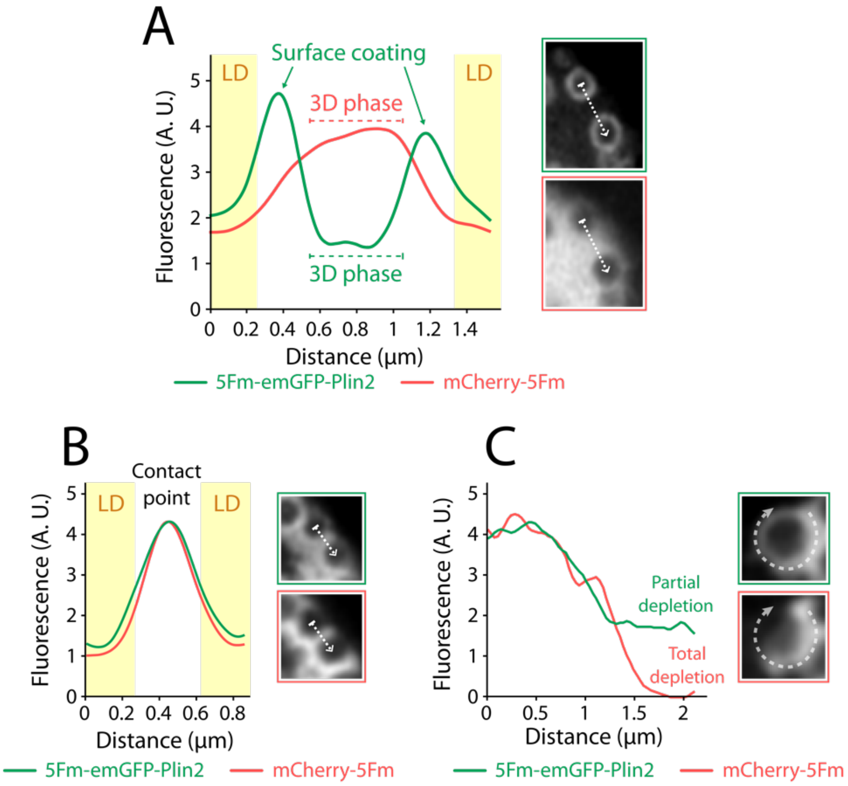


**Figure S1. Fluorescence profile in LD-condensates**

(A) Plot profile of fluorescence signals across two LDs within LD-condensates (extracted from Fig. 2A). Dashed arrow shows the profile section. The fluorescence background was set to the cytosol fluorescence signal, and the intensity scale is arbitrary. Yellow rectangles indicate LDs. (B.) Plot profile of fluorescence signals between two LDs within LD-condensates (extracted from Fig. 2C). The dashed arrow shows the profile location. The fluorescence background was set to the cytosol fluorescence signal, and the scale is arbitrary. Yellow rectangles indicate LDs. (C) Plot profile of fluorescence signals around an LD (extracted from Fig. 2C). The dashed arrow shows the profile location. The fluorescence background was set to the cytosol fluorescence signal, and the scale is arbitrary.

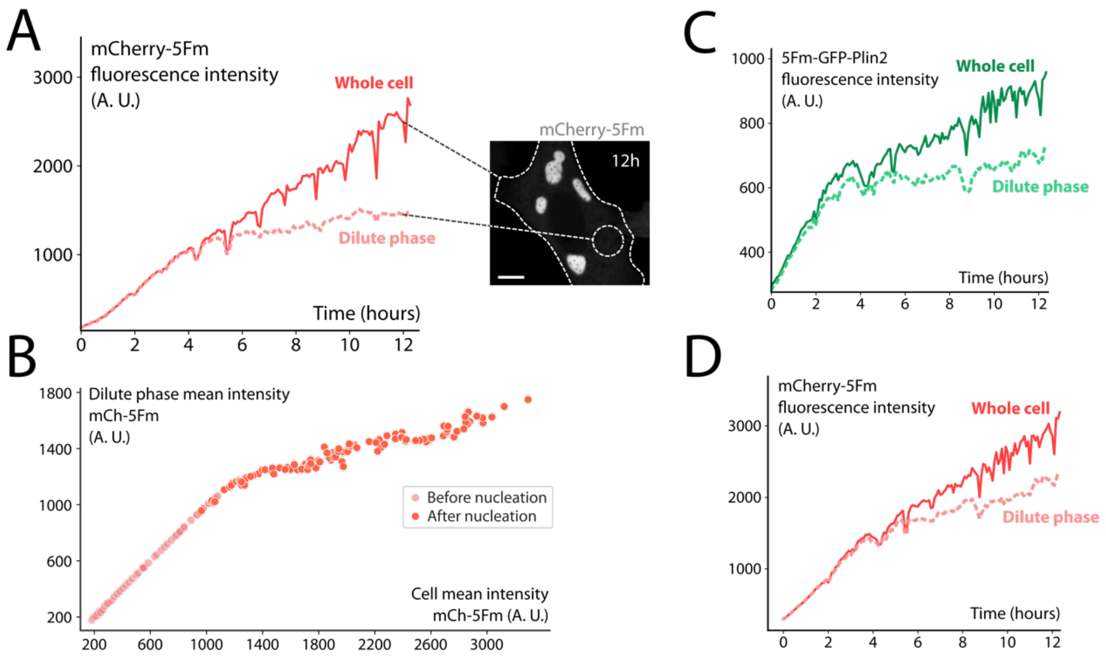


**Figure S2. Fluorescence intensities during LD-Condensates formation**

(A) mCherry mean fluorescence intensities during the formation of LD-condensates in the same HeLa cell as Fig. 3C. The whole cell intensity comprises the entire cell area, whereas the dilute phase intensity corresponds to the mean intensity of the areas outside LD-condensates. The event of nucleation correlates with the slope break of the dilute phase intensity. (B) Evolution of solubilized mCherry-5Fm fluorescence compared to total fluorescence, before and after condensates nucleation. (C) GFP mean fluorescence intensities during the formation of LD-condensates in another HeLa cell. (D) mCherry mean fluorescence intensities during the formation of LD-condensates in the same HeLa cell as Fig. S2B.

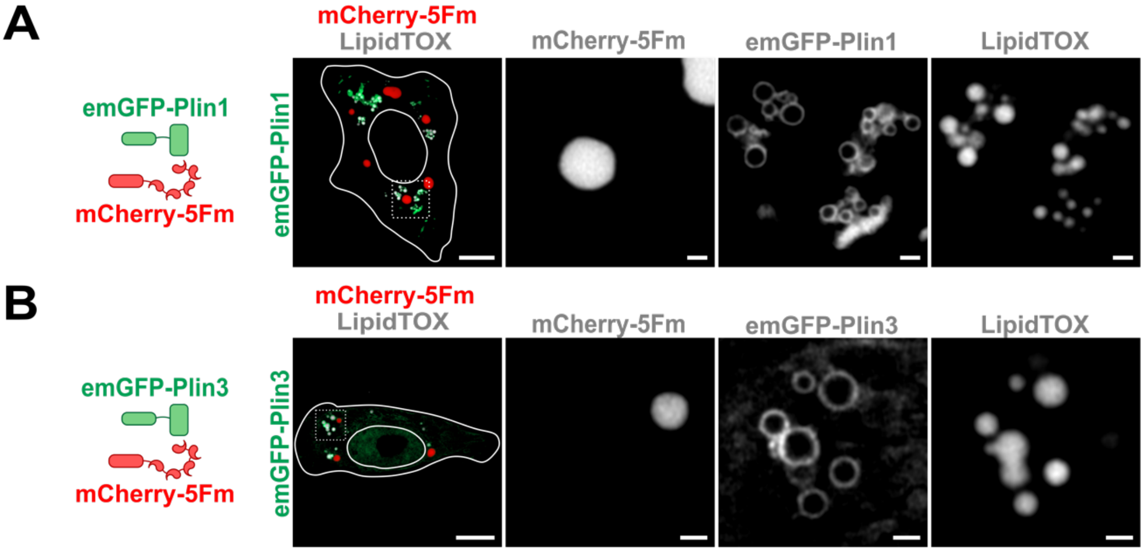


**Figure S3. Plin1 and Plin3 LD with 5Fm condensates**

(A) Airyscan confocal imaging of mCherry-5Fm condensates and Plin1-coated LDs in HeLa cells. Cells were fixed 24h after transfection with emGFP-Plin1, mCherry-5Fm, and 5Fm (plasmid ratio 1:5:14). LDs were marked using LipidTOX™ HCS. Grayscale images correspond to separate channels of the region of interest. Scale bar, 10 µm, 1 µm for zooms.
(B) Airyscan confocal imaging of mCherry-5Fm condensates and Plin3-coated LDs in HeLa cells. Cells were fixed 24h after transfection with emGFP-Plin3, mCherry-5Fm, and 5Fm (plasmid ratio 1:1:2). LDs were marked using LipidTOX™ HCS. Grayscale images correspond to separate channels of the region of interest. Scale bar, 10 µm, 1 µm for zooms.

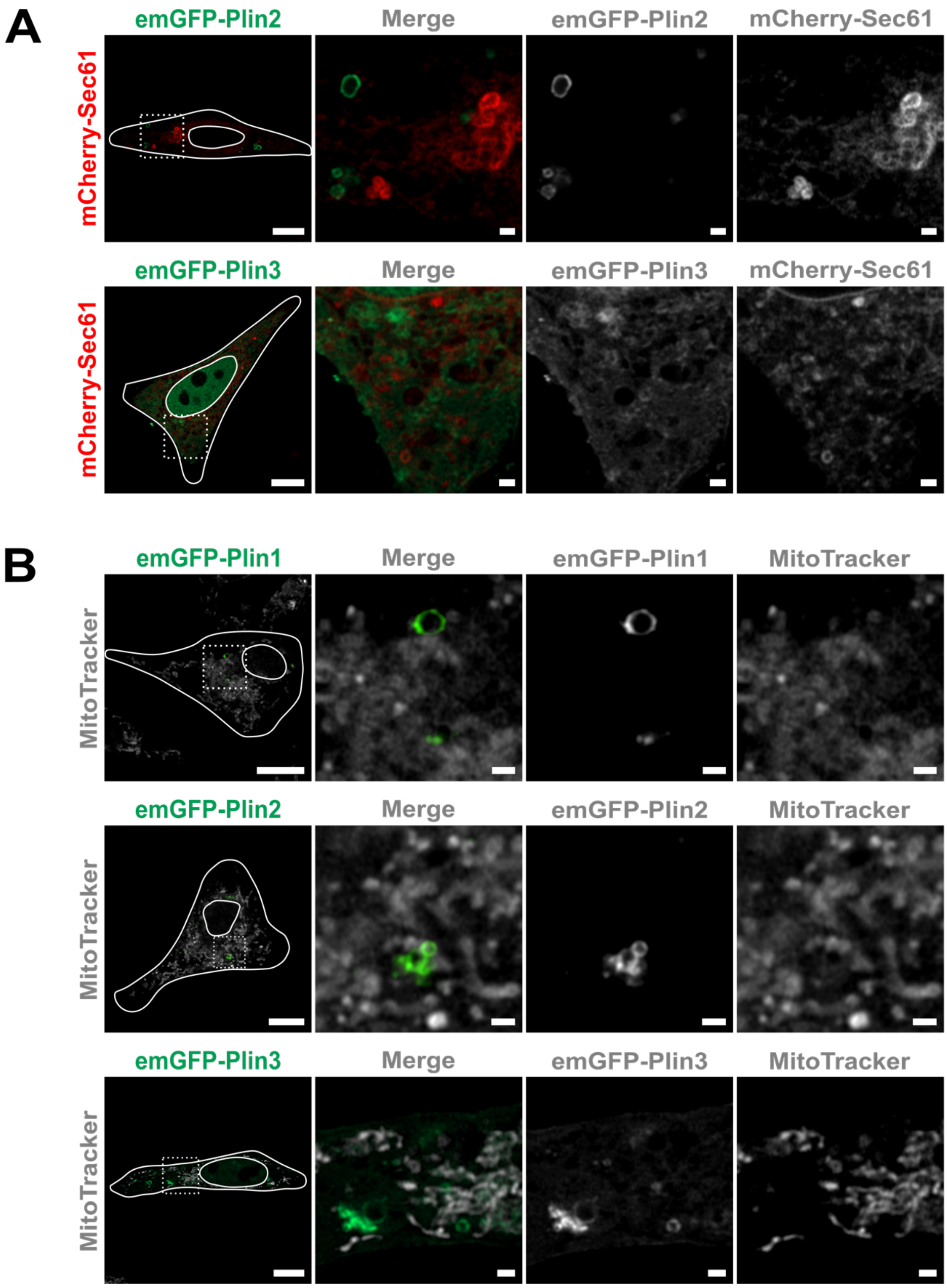


**Figure S4. Cells expressing Plin proteins with ER or mitochondria organelles markers**
(A) Airyscan confocal imaging of Plin2 or Plin3 in HeLa cells, and ER. Cells were fixed 24h after transfection with emGFP-Plin2 and mCherry-Sec61 or emGFP-Plin3 and mCherry-Sec61. Scale bar, 10 µm, 1 µm for zooms. (B) Airyscan confocal imaging of Plin1, Plin2 or Plin3 in HeLa cells, and mitochondria. Cells were fixed 24h after transfection with emGFP-Plin1 and mCherry-Sec61.

**Legend for Supplementary Movies**

**Supplementary Movie 1**: Time-lapse confocal imaging of the formation of LD-condensates in HeLa cells. Cells were transfected with 5Fm-emGFP-Plin2, mCherry-5Fm, and 5Fm (plasmid ratio 1:5:14) 6 hours before the start of the time-lapse. Images were taken every 5 min. Scale bar, 10 μm.

**Supplementary Movie 2**: Time-lapse epifluorescence imaging of the drug-induced dissolution of LD-trapping condensates in HeLa cells. Cells were transfected 24 hours before with 5Fm-emGFP-Plin2, mCherry-5Fm, and 5Fm (plasmid ratio 1:5:14). Dissolution is induced by exposition to FK506 (2.5 μM) at the start of the time-lapse, without mixing. Images were taken every 5 sec. Scale bar, 10 μm.
